## Supplementary Material for "Nuclear export is a limiting factor in eukaryotic mRNA metabolism"

### 5 Supplementary Material

#### 5.1 Basic read alignment

Reads were mapped to human reference genome hg19 with slamdunk [33] (version 0.3.0, settings: -n 100, -m) (Supplemental Figures S1, S2, S3 ). We chose slamdunk after comparison with bowtie2 [55] (version 2.2.6, default settings) and hisat2 [56] (version 2.0.0-beta, settings: -mp 1,1 , --no-spliced-alignment) (Supplemental Figure S1).

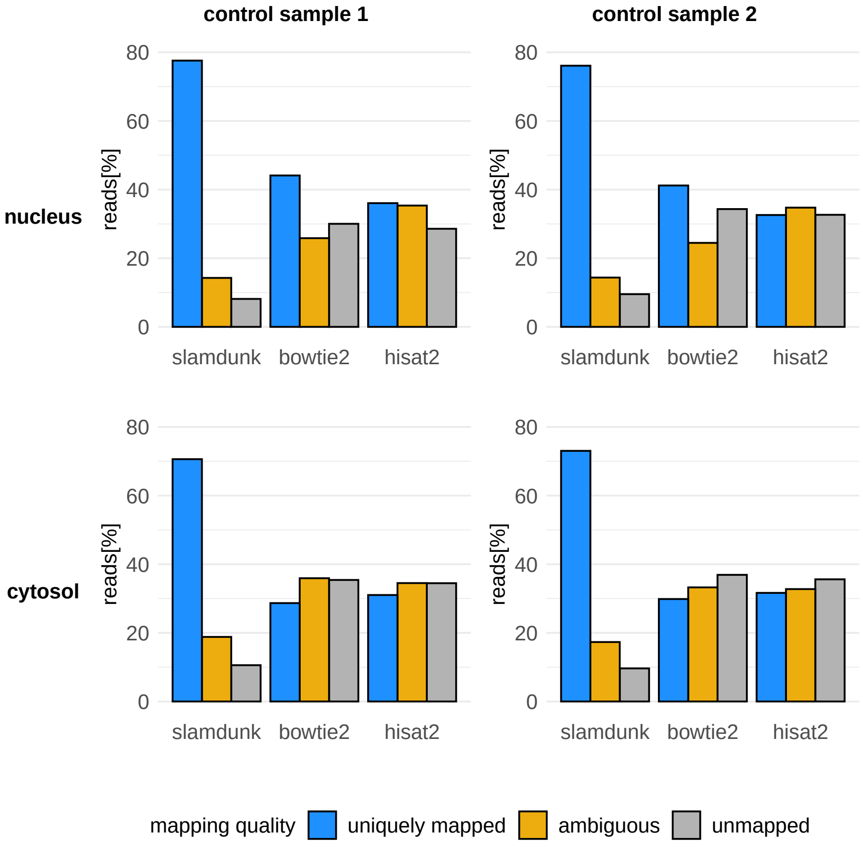


Supplemental Figure S1: Mapping statistics of the control data sets, shown for both control samples and the nuclear and cytosolic fraction. Uniquely mapped reads: reads with exactly one alignment (bowtie2 and hisat2), or reads with exactly one reliable alignment (slamdunk). Ambiguous reads: reads with more than one alignment (bowtie2 and hisat2), or reads with more than one reliable alignment or with only unreliable alignments (slamdunk). Unmapped reads: reads with no alignment (slamdunk, bowtie2, hisat2).

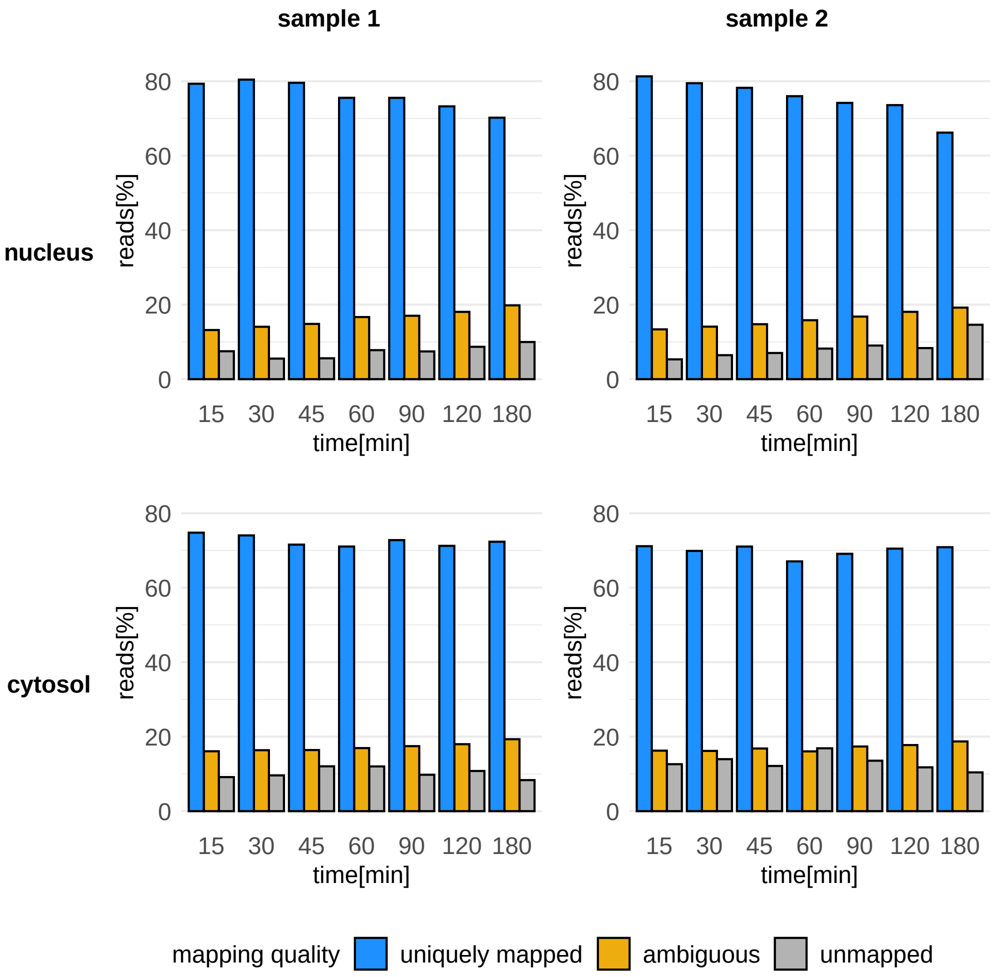


Supplemental Figure S2: Mapping statistics for both time series and the nuclear and cytosolic fraction. Uniquely mapped reads: reads with exactly one reliable alignment. Ambiguous reads: reads with more than one reliable alignment or with only unreliable alignments. Unmapped reads: reads with no alignment.

#### 5.2 Assignment of reads to 3'UTRs

The BED file defining hg19 annotated 3'UTRs was downloaded from UCSC [60]. Overlapping 3'UTRs were merged, considering strand orientation. Uniquely mapped reads were assigned to a specific 3'UTR if their alignment overlapped with the UTR with at least one nucleotide position. Mapping orientation was taken into account during read assignment, as reads derived from (-) strand genomic regions map forwards and reads derived from (+) strand genomic regions map reversely as a consequence of the experimental setup (Supplemental figure S3).

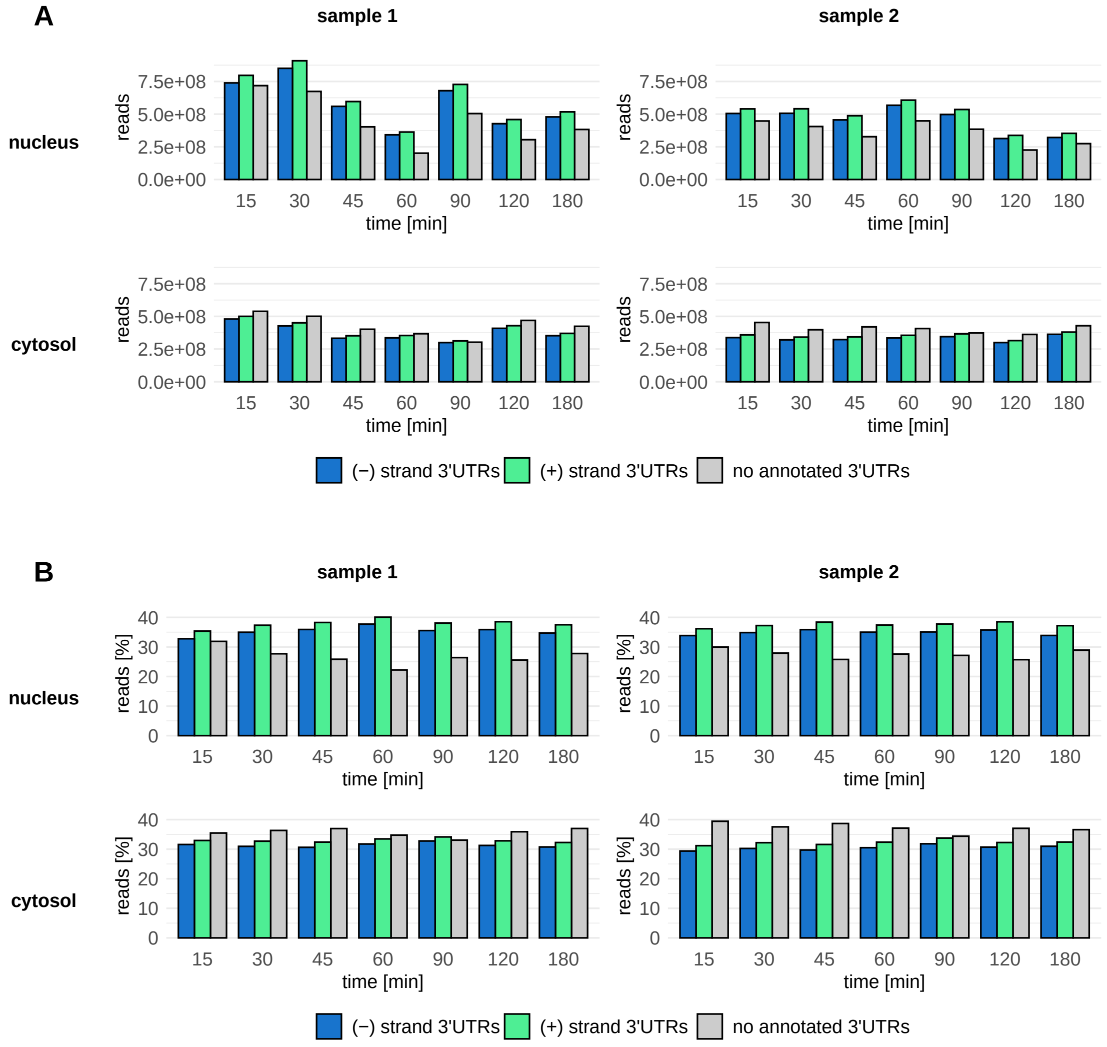


Supplemental Figure S3: Assignment of uniquely mapped reads of the SLAM-seq labeling samples to hg19 annotated 3'UTRs. Overlapping 3'UTRs were previously merged considering strand orientation. (A) Absolute read counts. (B) Relative read counts.

#### 5.3 Peak calling

To identify 3' transcript isoforms and transcripts with internal priming sites, we searched the genome for positions with a particularly high coverage of 3' ends of mapped reads ("peaks", to be defined below). By assigning only count to the 3' end position of a mapped read, we generated coverage profiles for the nuclear and cytosolic fractions of every measurement time point and each of the two replicate time series. Due biological variation and in the priming of the reverse transcription, the 3' poly-A site of a transcript might not be determined up to single nucleotide resolution. To obtain an (unbiased) idea on how spatially constrained such poly-A sites are in practice, we first binarized the coverage profiles in a ±1000bp window around each genomic position (0 = no count, 1 = at least one count at a given nucleotide position) and then averaged these profiles across all genomic positions that were covered at least once (Supplemental figures S4 and S5). The shape of these profiles indicated that counts typically aggregate in a $\pm 50$bp region around a central position, the closer the more.We therefore decided to define peak centers as the local coverage maxima of the genome-wide coverage profile that has been convolved with a triangular smoothing kernel of with $\pm 50$ (i.e., the function $f(x) = \max(1 - |x|/51, 0)$) (Supplemental figures S4 and S5). Peaks or peak regions are then the regions spanning -50bp to +50bp around a peak center.We identified a total of 98,102 peaks, 15,125 of which were measured with at least one count in each time series, time point and compartment, and 1,262,263 peaks for the reads not assigned to annotated 3'UTRs, 4481 of which were measured with at least one count in each time series, time point and compartment.

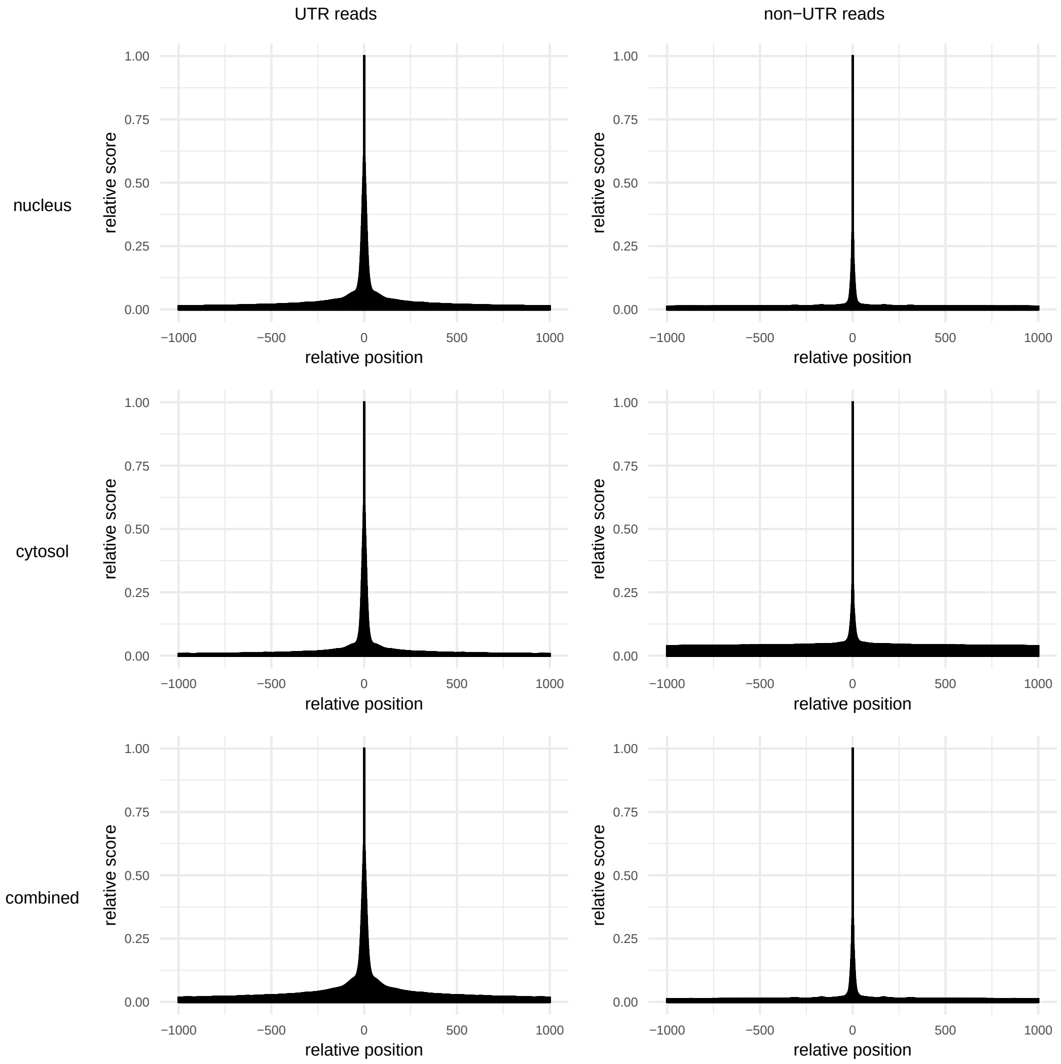


Supplemental Figure S4: Relative coverage profiles. Profiles were created separately for the reads assigned to annotated 3'UTRs (UTR-reads) and reads not assigned to such (non-UTR reads) as well as the nuclear and cytosolic fraction. The combined profile is based on the coverage of the nuclear and cyosolic fraction added up.

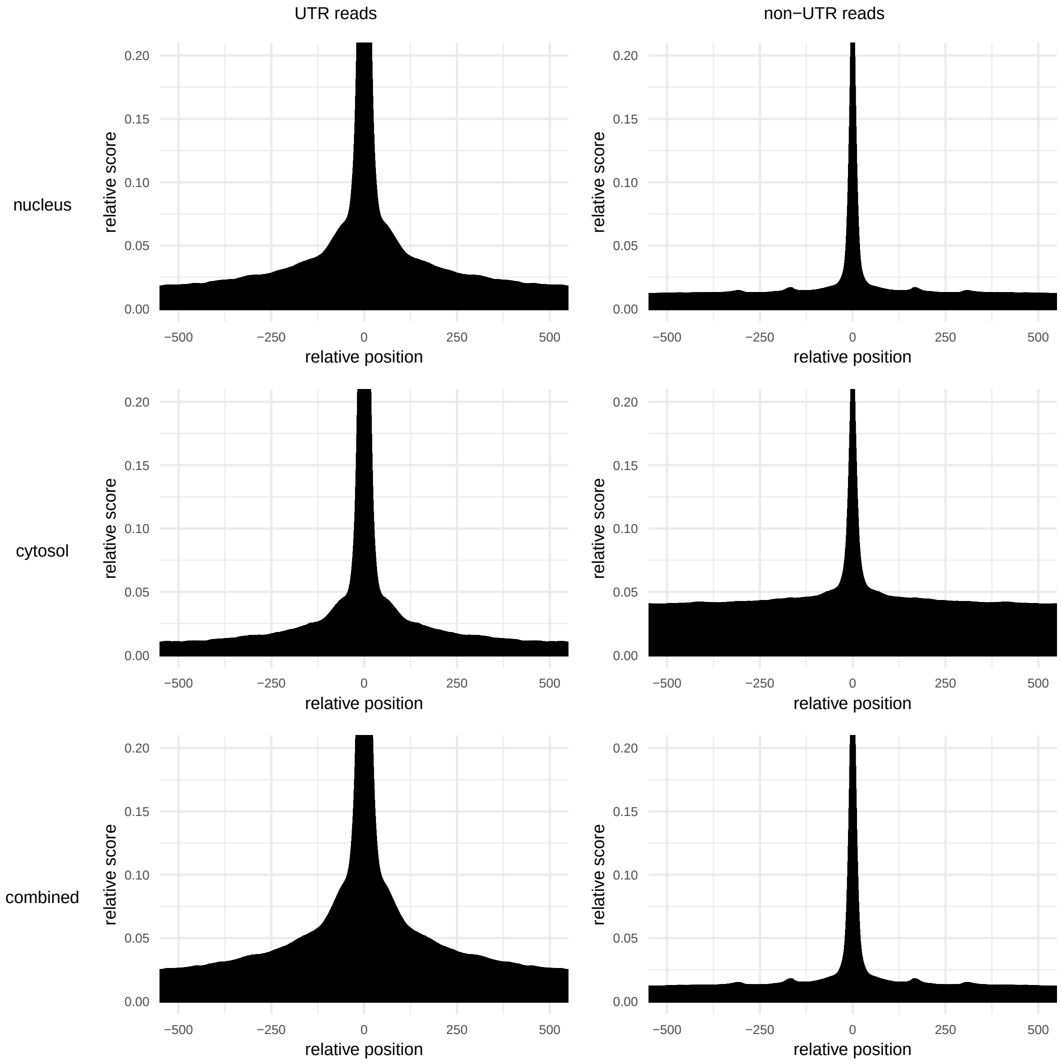


Supplemental Figure S5: Relative coverage profiles as shown in Supplemental figure S4 with trimmed y-axis for better visualization.

#### 5.4 Estimation of the labeling efficiency and new/total ratios

Recall the notation from the Methods. Fix a 3'UTR respectively a peak region onto which $J$ reads were mapped. Given a read $j = 1, ..., J$, let $T_j$ be its number of T-positions in the genomic sequence of the read alignment, and let $o_j \in 0, 1, ..., T_j$ be the number of positions at which we observe T>C transitions. Let $h_j \in \{0, 1\}$ be a hidden variable indicating whether read $j$ originates from a pre-existing RNA ($h_j = 0$), or a newly synthesized RNA ($h_j = 1$). Let $\rho \in [0, 1]$ be the proportion of newly synthesized RNAs in the RNA population from which read $j$ was drawn. Let $\ell \in [0, 1]$ be the labeling efficiency, i.e., the probability by which a T>C conversion happens in a newly synthesized RNA. Let $p_0 \in [0, 1]$ be the probability of a T>C sequencing error. Let $\epsilon \in [0, 1]$ be the probability of a C>non-C sequencing error, and denote by

$$p_1 = \ell(1 - \epsilon) + (1 - \ell)p_0 = \ell(1 - \epsilon - p_0) + p_0 \tag{9}$$

the probability of observing a T>C conversion in a newly synthesized read. Denote the parameters by $\Theta = (\rho, \ell, p_0, \epsilon)$.

To estimate sequencing error rates $p_0$ and $\epsilon$, we calculated the single-nucleotide mismatch rates for the nuclear and the cytosolic compartment in the control samples. We observed that, even after removal of SNP positions and A>I editing sites, the mismatch rates for the reads assigned to 3'UTRs showed an elevated T>C exchanges rate when mapping to the (+) strand and an elevated A>G exchange rate on the (-) strand (Supplemental Figure S30). We hypothesize this might be due to spurious A>I editing. The T>C probability $p_0$ was estimated as the average of the T>C respectively A>G exchange rate of reads mapping to the (+) respectivley (-) strand. The C>non-C probability $\epsilon$ was calculated as the average of the according, strand-specific base exchange rates.

From Equations (7,8), we have

$$\begin{aligned} \log P(o_j, h_j; \Theta) \quad &= \quad h_i \log \rho + (1-h_j)\log(1-\rho) + \log \binom{T_j}{o_j} \\ &\quad\; + o_j \log p_{h_j} + (T_j - o_j)\log(1-p_{h_j}) \end{aligned} \tag{10}$$

We employ an EM algorithm to estimate the ratio of newly synthesized RNAs $\rho$ and the labeling efficientcy $\ell$ for each 3'UTR separately for each compartment and time point in each time series. To that end, we initiliased the labeliung efficiency l with $p_0$ and the $\frac{new}{total}$ RNA ratio $\rho$with the observed $\frac{labeled}{total}$ RNA ratio.

**E-step.** Let $H = (h_j; j = 1, ..., J)$, $H_{-1} = (h_j; j = 2, ..., J)$. Given some parameter set $\Theta' = (\rho'$,e'), we have to optimize the lower bound function $Q(\Theta; \Theta')$ with respect to $\Theta = (\rho, e)$. Using standard algebraic transformations of the lower bound function, we obtain

$$\begin{aligned} Q(\Theta;\Theta') \quad &:= \quad \mathbb{E}_{P(H|O;\Theta')} \log P(O, H; \Theta) \\ &= \quad \sum_{H \in \{0,1\}^J} P(H \mid O; \Theta') \log P(O, H; \Theta) \\ &= \quad \sum_{H_{-1} = (h_2,...,h_J) \in \{0,1\}^{J-1}} \; \sum_{h_1 \in \{0,1\}} (P(H_{-1} \mid O_{-1}; \Theta') \cdot P(o_1, h_1; \Theta')) \\ &\quad\quad \cdot (\log P(H_{-1} \mid O_{-1}; \Theta) + \log P(o_1, h_1; \Theta)) \\ &= \quad \sum_{h_1 \in \{0,1\}} P(h_1 \mid o_1; \Theta') \log P(o_1, h_1; \Theta) \\ &\quad\quad + \sum_{H_{-1} \in \{0,1\}^{J-1}} P(H_{-1} \mid O_{-1}; \Theta') \log P(O_{-1}, H_{-1}; \Theta) \\ &\overset{\text{induction}}{=} \quad \sum_{j=1}^{J} \sum_{h_j \in \{0,1\}} P(h_j \mid o_j; \Theta') \log P(o_j, h_j; \Theta) \end{aligned} \tag{11}$$

Let

$$c_{j,h_j} := P(h_j \mid o_j; \Theta') = \frac{P(o_j \mid h_j; \Theta') P(h_j; \Theta')}{\sum_{h_j \in \{0,1\}} P(o_j \mid h_j; \Theta') P(h_j; \Theta')} \quad , \; j = 1, ..., J \tag{12}$$

and $C_0 = \sum_j c_{j,0}$ and $C_1 = \sum_j c_{j,1}$, $A = \sum_j c_{j,1} o_j$, $B = \sum_j c_{j,1}(T_j - o_j)$. Then, (11) simplifies to

$$\begin{aligned} Q(\Theta;\Theta') &= \sum_j c_{j,0}\left[\log(1-\rho) + \log\binom{T_j}{o_j} + o_j \log p_0 + (T_j - o_j)\log(1-p_0)\right] \\ &\quad + \sum_j c_{j,1}\left[\log\rho + \log\binom{T_j}{o_j} + o_j \log p_1 + (T_j - o_j)\log(1-p_1)\right] \\ &= C_0\log(1-\rho) + C_1\log\rho + A\log p_1 + B\log(1-p_1) + \text{const} \end{aligned} \tag{13}$$

where all terms that do not depend on $\ell$ or $\rho$ are summarized in the constant.

**M-step.** Taking the partial derivative of $Q$ with respect to $\rho$ and equating this expression to zero yields

$$\begin{aligned} 0 = \frac{\partial Q(\Theta;\Theta')}{\partial \rho} &= -\frac{C_0}{1-\rho} + \frac{C_1}{\rho} \\ \rho &= \frac{C_1}{C_0 + C_1} = \frac{C_1}{J} \end{aligned} \tag{14}$$

Equating the partial derivative of $Q$ with respect to $\ell$ to zero yields

$$\begin{aligned} 0 = \frac{\partial Q(\Theta;\Theta')}{\partial \ell} &= \frac{A}{p_1} \cdot \frac{\partial p_1}{\partial \ell} + \frac{B}{1-p_1} \cdot \frac{\partial(1-p_1)}{\partial \ell} \\ &= \frac{A}{p_1} \cdot (1-\epsilon-p_0) - \frac{B}{1-p_1} \cdot (1-\epsilon-p_0) \end{aligned} \tag{15}$$

Solving this for $p_1$ (assuming $1-\epsilon-p_0 \neq 0$) and then solving Equation (9) for $\ell$ yields

$$\begin{aligned} p_1 &= \frac{B}{A+B} \\ \ell &= \frac{p_1 - p_0}{1-\epsilon-p_0} \end{aligned} \tag{16}$$

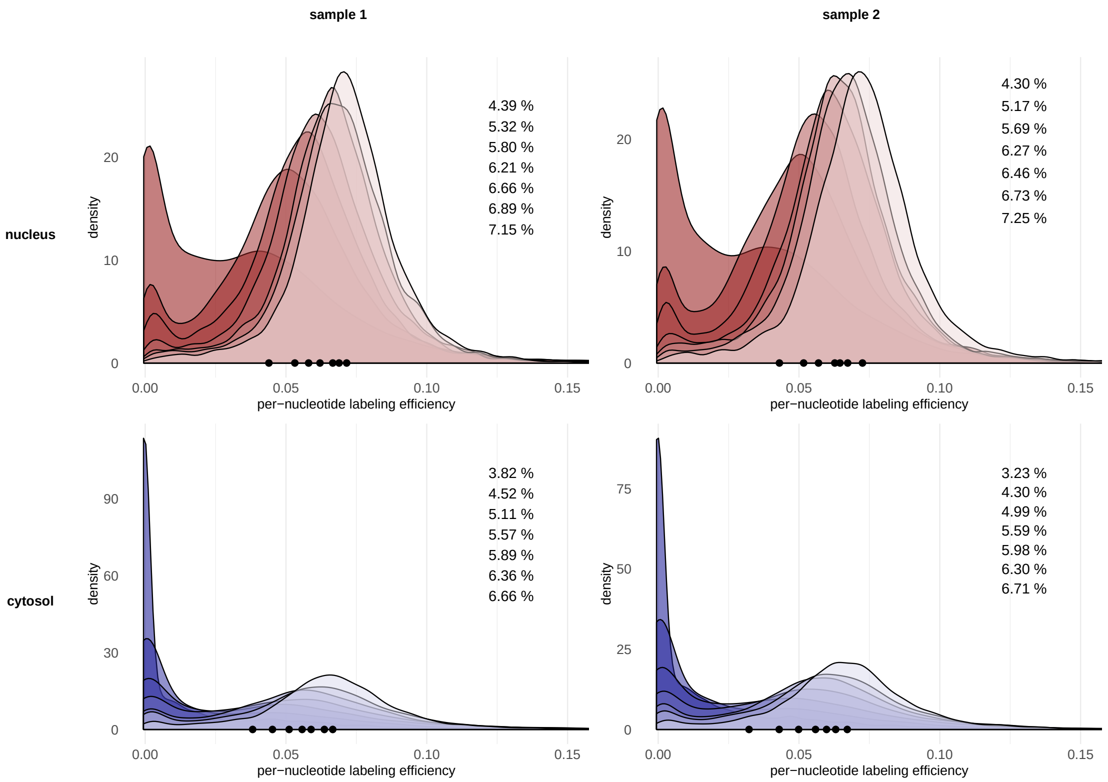


Supplemental Figure S6: Distribution of labeling efficiency estimates $\ell$, calculated for each 3'UTR and each compartment, time point and time series. Each distribution represents one time point for the respective time series and compartment, with darker shades of a color encoding earlier and brighter shades encoding later time points. Efficiencies were estimated for all 3'UTRs with a coverage of at least 100 in the respective measurement. The black dots and text labels indicate the median labeling efficiencies obtained after discarding all 3'UTRs with an efficiency estimate smaller than 0.01 within a measurement. Text labels are ordered by increasing measurement time points. The x-axis is trimmed for better display.

#### 5.5 Modeling of the labeling efficiency in the nucleus and the maturation gap

**General considerations on the labeling efficiency.** The parameters of our model, such as RNA synthesis or RNA degradation, were so far assumed to be constant. While we have chosen minimally invasive procedures to keep these parameters unchanged, the assumption of constant labeling efficiency can be problematic, as discussed below. To understand this, remember that the time variable $t$, which appears in our dynamic equations, denotes the time after the start of 4sU labeling. When we say that a transcript has been synthesized at time $t$, we mean that this transcript has been polyadenylated and can be detected with our experimental method at time $t$. Let $l(t)$ be the labeling efficiency at time $t$, i.e., the probability that a 4sU nucleotide instead of uridine is included in a nascent transcript at time $t$. Because it may take a while for the 4sU molecules to diffuse into the nucleus, we expect the labeling efficiency to increase monotonically from $l(0) = 0$ to a certain limit.

First, we verify the assumption implicit in our model that labeling efficiency is the same for each uridine position of the same transcript. The length of a primary human gene transcript (including introns) ranges between 0.5kb and 1,000kb [61]. As the elongation speed of RNA Polymerase II ranges between 0.5kb/min and 5kb/min[62], the synthesis of a primary transcript can take up to a few hours. The onset of transcription may therefore even precede the start of 4sU labeling in long genes, which seems inconsistent with the assumption of constant labeling efficiency. However, this apparent contradiction resolves if we considered that we are sequencing only the 3' ends of mRNAs, i.e., the last ˜80nt upstream of the poly(A) tail of a transcript. All considerations on labeling efficiency refer only to this 3' end, which is synthesized by RNA polymerase II within seconds. It therefore seems justified to assume that the labeling efficiency during 3'-end synthesis of a transcript is constant and equals $l(t)$. Note that we still have neglected the time it takes until the poly(A) tail has been added after transcription termination. As a consequence, we will observe all processes described above with a delay. We call this delay the maturation gap $\Delta$, which we expect to be in the range of seconds to minutes.

**Estimation of the labeling efficiency kinetics.** Denote by $\mathrm{RNA}_x^{new}(t)$ the amount of (mature) RNA of transcript $x$ in the nucleus that has been newly synthesized (i.e., is detecteble by RNA-seq) within $t$ minutes after the start of 4sU labeling. Let $\mathrm{RNA}_x^{lab}(t)$ the amount of newly synthesized RNA that can be detected as labeled (by at least one T>C transition) within $t$ minutes after labeling start. Our next goal is to estimate the fraction $f_x(t) = \mathrm{RNA}_x^{lab}(t)/\mathrm{RNA}_x^{new}(t)$ of newly synthesized RNA of a transcript $x$ at time $t$ which is recognized as labeled. This fraction will be used later as a correction factor in the estimation process. We assume that the labeling efficiency in the nucleus follows a delayed saturation kinetics,

$$l(t) = l(t) = \ell_{max} \cdot max(1 - e^{-\gamma(t-\Delta)}, 0) \tag{17}$$

Here, $\ell_{max}$ is the limit 4sU incorporation rate, $\gamma$ models the diffusion speed of $4sU$ inside the cell, and $\Delta$ is the maturation gap. Let $u_x > 1$ be the number of Uridines in the sequenced part of transcript $x$. The lengths of the sequenced reads are very similar, so without loss we may assume this number to be constant for each transcript of type $x$. Then, the probability of a transcript $x$ that has been newly synthesized exactly at time $t$ to be recognized as labeled is

$$r_x(t) = 1 - (1 - l(t))^{u_x} \tag{18}$$

Assuming a constant synthesis rate $\mu_x$ and a constant export+decay rate $\lambda_x$ for $x$, the ordinary differential equations for the nuclear RNA become

$$\begin{aligned} \frac{d}{dt}\mathrm{RNA}_x^{new}(t) &= \mu_x - \lambda_x \mathrm{RNA}_x^{new}(t) \qquad (19) \\ \frac{d}{dt}\mathrm{RNA}_x^{lab}(t) &= \mu_x \cdot r_x(t) - \lambda_x \mathrm{RNA}^{lab}(t) \end{aligned}$$

They have the solutions

$$\begin{aligned} \mathrm{RNA}_x^{new}(t) &= \mu_x \frac{1 - e^{-\lambda_x t}}{\lambda_x} \qquad (20) \\ \mathrm{RNA}_x^{lab}(t) &= \mu_x \int_0^t r_x(t') e^{\lambda_x (t'-t)} dt' \end{aligned}$$

Thus, the fraction $f_x(t)$ of newly synthesized RNAs that are detected as labeled/converted is

$$f_x(t) = \frac{\mathrm{RNA}_x^{lab}(t)}{\mathrm{RNA}_x^{new}(t)} = \frac{\lambda_x \cdot \int_0^t r_x(t')e^{\lambda_x(t'-t)}dt'}{1-e^{-\lambda_x t}} \tag{21}$$

It remains to estimate $s_{max}$ and $\gamma$. If $t'$ is the time after labeling start at which a transcript $x$ has been synthesized, the distribution of $t'$ in the population of newly synthesized transcripts $x$ at time $t$ after labeling is

$$A_x(t';t,\lambda_x) = e^{\lambda_x(t'-t)} / \int_0^t e^{\lambda_x(s-t)}ds = \lambda_x e^{\lambda_x t'}/(e^{\lambda_x t}-1) \quad , \; t' \in (0,t] \tag{22}$$

We conclude that the average labeling efficiency measured for a single uridine in a newly synthesized transcript $x$ at measurement time $t$ is

$$e_x(t;\lambda_x,\gamma,\ell_{max},\Delta) = \int_0^t l(t')A(t',t,\lambda_x)dt' \tag{23}$$

We have assessed how $e_x(t;\lambda_x,\gamma,\ell_{max})$ varies with transcript half lives, respectively their degradation rate $\lambda_x$ (Fig S8). We found that for fixed time points $t$, $e_x$ is approximately independent of $\lambda_x$, for $\lambda_x$ corresponding to transcript half lives above $\sim$ 100min (i.e., relative changes are small). The reason is that for sufficiently small $\lambda_x$, $A(t',t,\lambda_x)$ is approximately constant for all $t' \in (0,t]$ except for very small values of $t'$. We therefore define the (degradation-rate independent) labeling efficiency

$$\bar{e}(t;\gamma,\ell_{max},\Delta) := \int_0^t l(t')dt' = \ell_{max}\left(1 - \frac{1-e^{-\gamma(t-\Delta)}}{\gamma(t-\Delta)}\right) \tag{24}$$

and keep in mind that

$$\bar{e}(t;\gamma,\ell_{max},\Delta) \approx e_x(t;\lambda_x,\gamma,\ell_{max},\Delta) \tag{25}$$

for $\lambda_x$ small enough, say for a transcript half life greater than 100min.

In a first round of our estimation procedure, we assume that labeling efficiencies do not depend on the degradation rate of a transcript. Using our estimation procedure (Supplements 5.4), we obtain preliminary labeling efficiency estimates $\hat{e}_x(t)$ for each measurement time $t$ and transcript $x$, together with preliminary half life estimates for each transcript $x$. We found that 98% of all transcripts have half life estimates above 100min. Since the median is a robust measure of location for skewed distributions, we use the median of all $\hat{e}_x(t)$ for transcripts with a preliminary half life estimate >100min as an estimate of $\bar{e}(t;\gamma,\ell_{max})$:

$$\hat{e}(t) := \mathrm{median}\left(\hat{e}_x(t);\, \text{prelim. half life}\lambda_x > 100\text{min}\right) \tag{26}$$

We additionally account for the maturation gap by introducing a time shift of $\Delta$ for our observations. The estimates of $\gamma$, $s_{max}$ and $\Delta$ are then obtained by a least squares fit of

$$\hat{e}(t) \sim \bar{e}(t;\gamma,\ell_{max},\Delta) \quad , \; t \in \{15, 30, 45, 60, 90, 120, 180\text{min}\} \tag{27}$$

This yields (see SFig S7)

$$\ell_{max} = 7.8\% \ (7.9\%) \tag{28}$$
$$\gamma = 0.069\text{min}^{-1} \ (0.056\text{min}^{-1}) \tag{29}$$
$$\Delta = -14\text{min} \ (-18\text{min}) \tag{30}$$

for replicate time series 1 (replicate time series 2). Contrary to our intuition, the estimated value for the maturation gap was negative. We still keep our model, since it represents an excellent fit of labeling efficiency time curves.

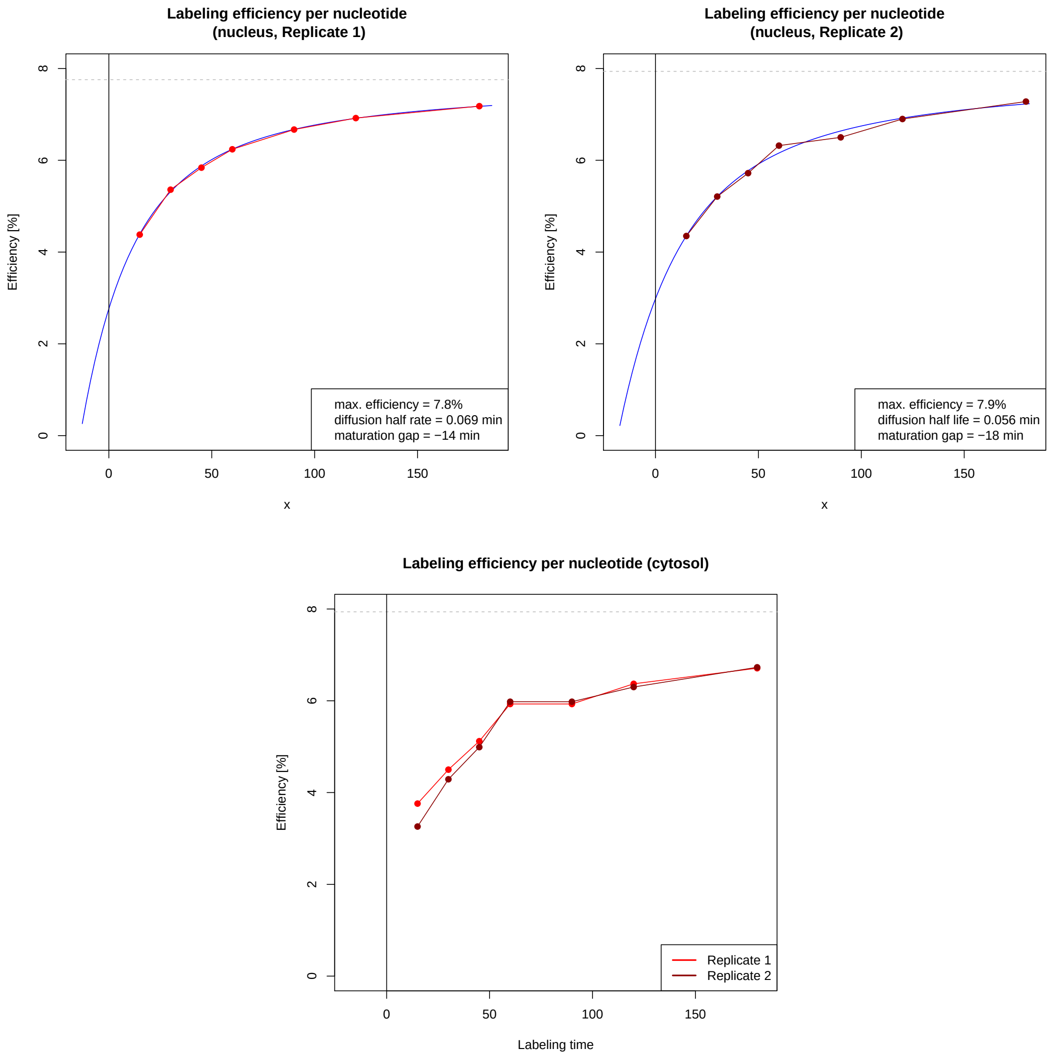


Supplemental Figure S7: Per nucleotide labeling efficiencies. The dots (red/dark red: replicate 1/replicate 2) show the median values for $\hat{e}(t)$ for each measurement time point (top: nucleus, bottom: cytosol). The blue lines are the least squares fits for the parameters $\gamma$, $\ell_{max}$ and $\Delta$ in both replicate time series. The parameters of the labeling efficiency function $\bar{e}(t-\Delta;\gamma,\ell_{max})$ (Equation (24)) were fitted according to Equation (27).

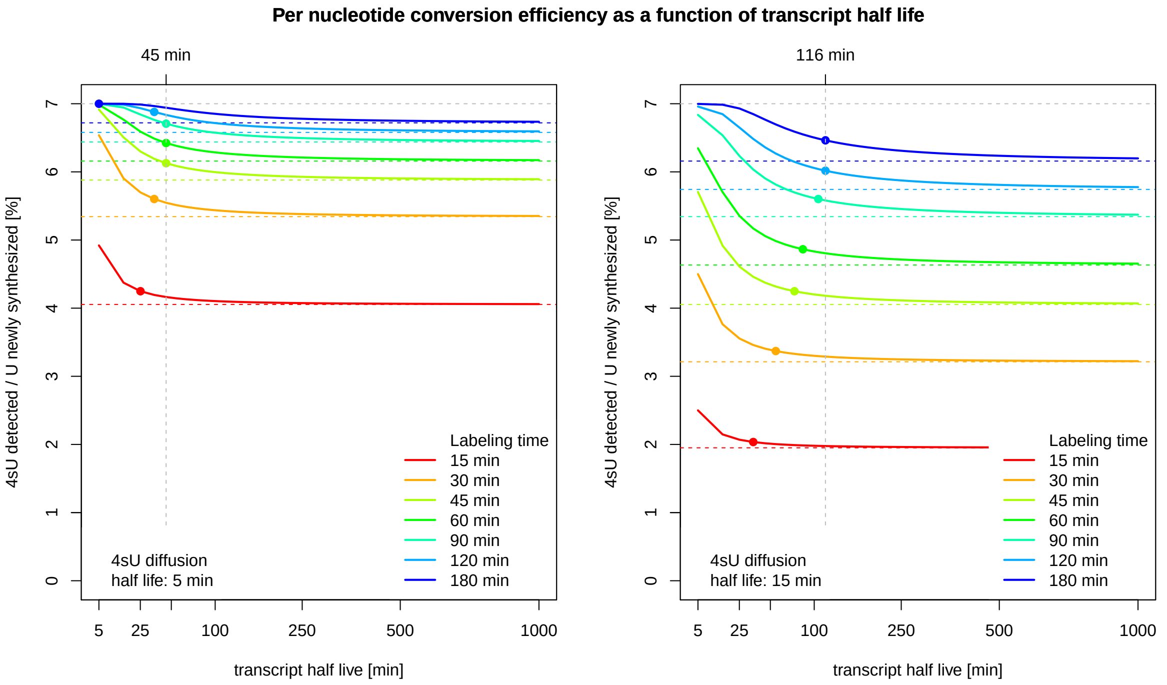


Supplemental Figure S8: Per nucleotide labeling+conversion efficiency ($e(t; \lambda_x, \gamma, \ell_{max}, g)$, see Equation (23)) as a function of transcript halflife ($\log(2)/\lambda_x$, x-axis). This was done for a diffusion rate $\gamma$ coresponding to a half live $(\log 2)/\gamma$ of 5min (left plot) and a half life of 15min (right plot), which is considered a realistic upper bound for $\gamma$. The maximum labeling efficiency was set to $\ell_{max} = 7\%$ (dashed gray horizontal lines). Each colored curve corresoponds to a certain labeling time $t = 15, 30, ..., 180$min (see the inset). The dashed colored horizontal lines indicate the corresponding values of $\bar{e}(t; \gamma, \ell_{max})$. The colored dots on the curves indicate the half lives above which Approximation (25) has an error smaller than 5%. The dashed gray vertial line at 45min (left) respectively 116min (right) indicates the transcript half life above which Approximation (25) has an error smaller than 5% for all measurement time points.

**Necessity of modeling time-dependency of the labeling efficiency.** So far, half life estimation procedures based on SLAM-Seq assumed a labeling efficiency that is constant over time [28]. This means replacing the above labeling efficiency estimate $f_x(t)$ by an approximation $g_x(t)$ assuming constant per-uridine labeling efficiency $\bar{e}_x(t)$:

$$g_x(t) = 1 - (1 - \bar{e}_x(t))^{u_x}$$

We have assessed whether this assumption causes a relevant bias, or whether this simplification is justified. Starting with the equation

$$\frac{\text{RNA}_x^{lab}(t)}{\text{RNA}_x^{total}(t)} = \frac{\text{RNA}_x^{lab}(t)}{\text{RNA}_x^{new}(t)} \cdot \frac{\text{RNA}_x^{new}(t)}{\text{RNA}_x^{total}(t)} = f_x(t) \cdot (1 - e^{-\lambda_x t})$$

we replace $f_x(t)$ by $g_x(t)$ in the estimation process. This amounts to letting

$$g_x(t) \cdot (1 - e^{-\hat{\lambda}_x t}) \approx \frac{\text{RNA}_x^{lab}(t)}{\text{RNA}_x^{total}(t)} = f_x(t) \cdot (1 - e^{-\lambda_x t})$$

If we merely use the observation at measurement time point $t$, the biased estimate $\hat{\lambda}_{x,t}$ of $\lambda_x$ becomes

$$\hat{\lambda}_{x,t} = -\frac{1}{t} \log \left[ 1 - \frac{f_x(t)}{g_x(t)} \cdot \left(1 - e^{-\lambda_x t}\right) \right]$$

In a post-hoc analysis, we fix the values $\hat{\gamma}$ and $\hat{s}_{max}$ to the values obtained from replicate time series 1, see Equations (28,28). We then calculated the the relative bias $\hat{\lambda}_{x,t}/\lambda_x$ for all our measurement time points $t$, for the whole range of transcript half lives $\lambda_x$, and for various uridine contents of the reads (SFig S9). The relative bias is generally negative for short half lives (except for very high uridine contents and very short labling times). The longer the labeling times, the more the bias is shifted towards negative values. Above a half life of ˜100min, the bias is asymptotically constant as a function of half life, and it ranges in the interval $[-7\%, 7\%]$, with longer labeling times resulting in smaller biases. In conclusion, the estimation bias relatively small for the setting in our experiment. We therefore do correct for the time-dependency of the labeling efficiency, but we generally advise to perform an assessment of this bias in future SLAM-Seq experiments.

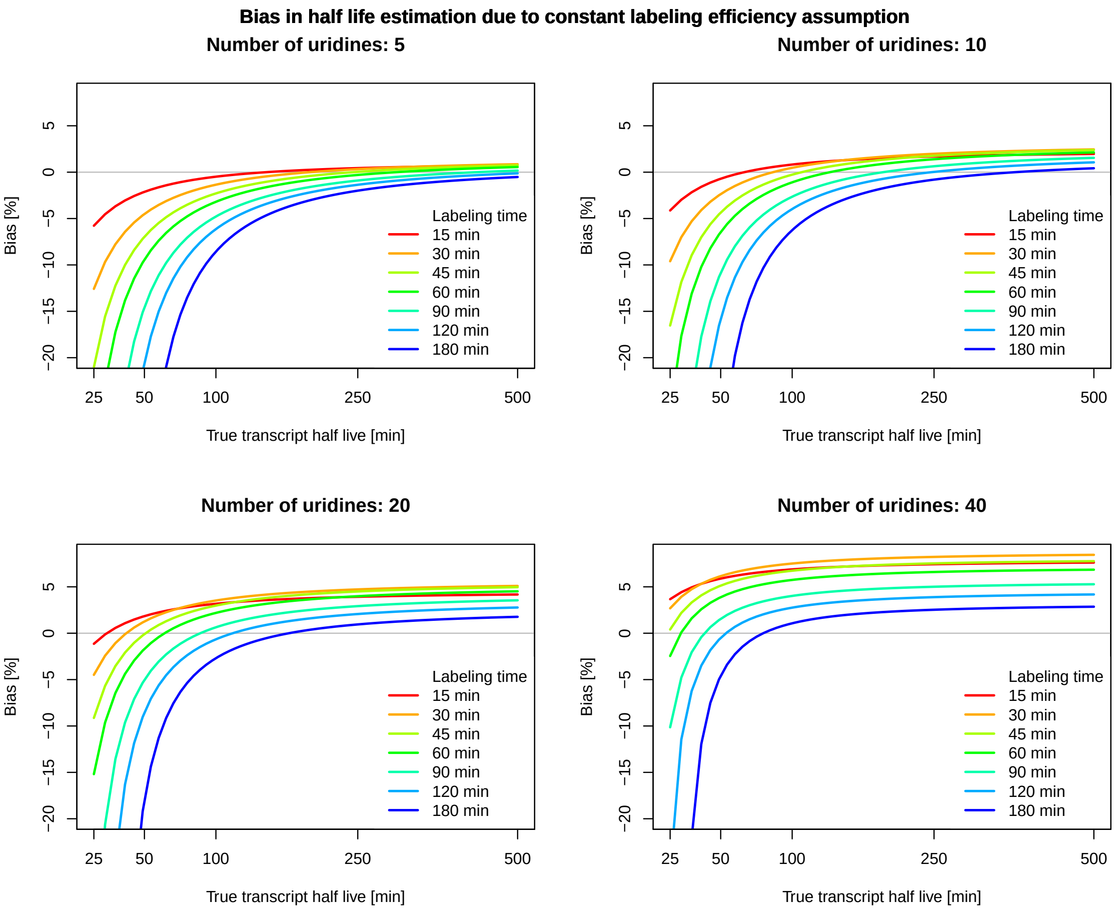


Supplemental Figure S9: Assessment of half life estimation bias when assuming constant labeling efficiency. Each line shows the relative bias (in %) as a function of the (true) transcript half life. Line colors correspond to different labeling times (see insets). The caclulations were done for different uridine contents of a transcript: 5 (top left), 10 (top right), 20 (bottom left), 40 (bottom right) uridines per sequenced read. The time dependency of the labeling efficiency was modeled by Equation (17) with parameters $\gamma = 0.069\text{min}^{-1}$ and $\ell_{max} = 7.8\%$ estimated from replicate time series 1.

#### 5.6 Effects of variance stabilization

We thoroughly assessed the effects of variance stabilization of the estimates $\frac{new}{total}$RNA ratios as described in Methods 4.5 (Supplemental Figure S10)

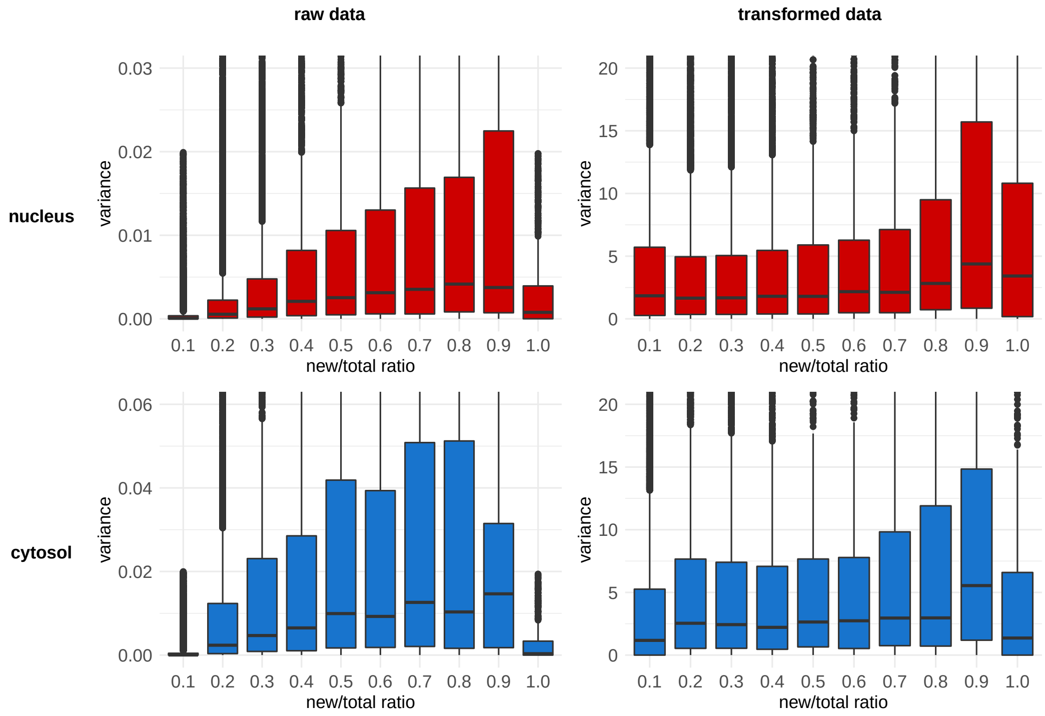


Supplemental Figure S10: Left: Variance of new/total mRNA ratios (y-axis) as a function of the mean (x-axis, binned in steps of 0.1), for the nuclear (red) and cytosolic (blue) fraction. Variance was calculated as $\frac{1}{2}(\rho_1 - \rho_2)^2$, where $\rho_i$ are the estimated new/total ratios from the two replicate time series, for all time points and all transcripts. Right: Same as left, with $\rho_i$ replaced by $\sqrt{4N_i}\text{arcsin}\sqrt{\rho_i}$.

#### 5.7 Assessment of parameter estimate variability

To check the variability in our parameter estimates, we correlated the half-lives of both time series replicates. This was done seperately for the nuclear and cytosolic half-life estimates (Supplementary Figure S11-S12).

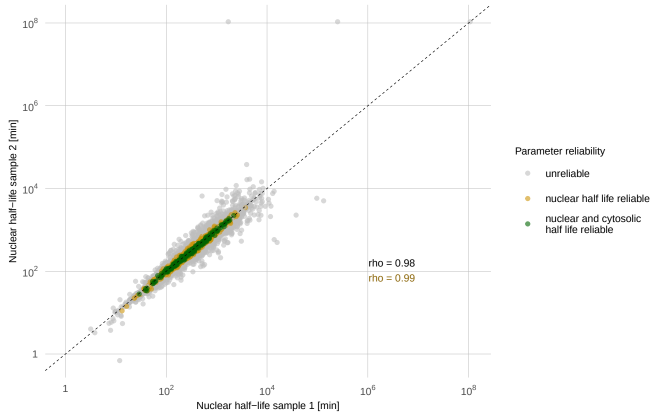


Supplemental Figure S11: Correlation between nuclear half-lives obtained from two distinct SLAM-seq time-series.

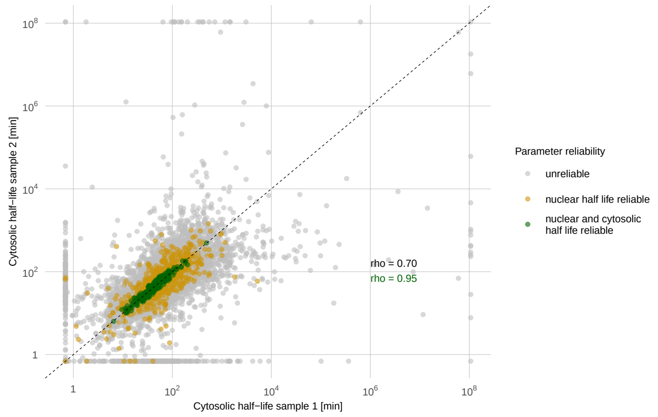


Supplemental Figure S12: Correlation between cytosolic half-lives obtained from two distinct SLAM-seq time-series.

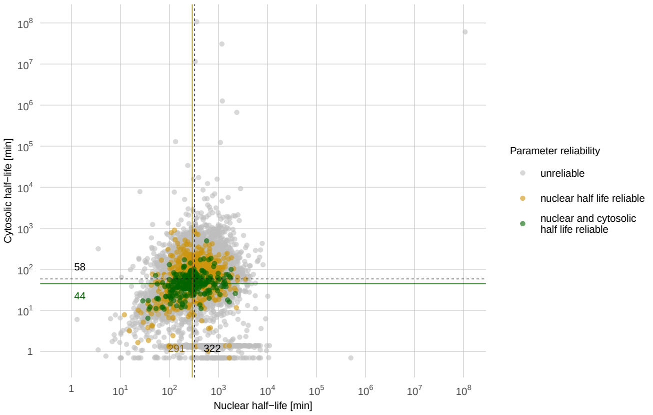


Supplemental Figure S13: Nuclear and cytosolic RNA half-life estimates of 3'UTRs. Half-lives were averaged over both measured time series. Stringent quality criteria were applied to score the reliability of the estimates. gray dots represent unreliable estimates, yellow dots correspond to estimates that passed additional reliability criteria in the nucleus, and green dots portray estimates that matched the reliability criteria in the nuclear and cytosolic compartment.

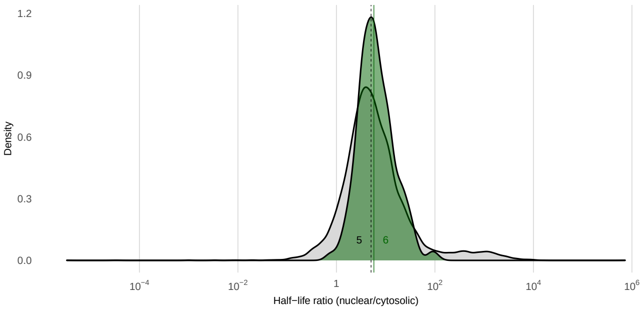


Supplemental Figure S14: Nuclear by cytosolic half-life ratios of all 3'UTRs and 3'UTRs with reliable half-life estimates for both compartments. The solid lines indicate median half-lives.

#### 5.8 Comparison with half-life estimates from literature

The compartment-specific parameter estimates for a certain transcript can be converted into an estimate of its RNA half life / degradation rate in the entire cell. Let $N_\infty = \frac{\mu}{\tau+\nu}$ and $C_\infty = N_\infty \cdot \frac{\tau}{\lambda}$ the RNA abundances of a certain transcript in the nucleus respectively cytosol. Assuming a one-compartment model with total steady-state RNA abundance $N_\infty + C_\infty$, a constant synthesis rate $\mu$ and degradation rate $\lambda_w$, the steady-state relation is

$$\frac{\mu}{\lambda_w} = N_\infty + C_\infty = \mu \left( \frac{1}{\tau + \nu} + \frac{1}{\lambda} \cdot \frac{\tau}{\tau + \nu} \right)$$

Assuming $\nu \approx 0$, the whole-cell half life $h_w$ is simply the sum of the nuclear half life $h_n$ and the cytosolic half life $h_c$:

$$h_w := \frac{\log 2}{\lambda_w} = \frac{\log 2}{\nu + \tau} + \frac{\log 2}{\lambda} \cdot \frac{\tau}{\nu + \tau} \approx h_n + h_c$$

This surrogate half life $h_w$ is used to compare our results with half life estimates from the literature.

For comparison of our RNA half-lives with estimates from the literature, we summarized our nuclear and cytosolic half-life estimates. These pseudo-WCE half-lives where correlated against estimates from Schueler et al. (2014) [37] (Figure 2C,Supplementary Figure S15). To that end, we averaged their replicates for both the MCF7 and HEK293 cell lines.

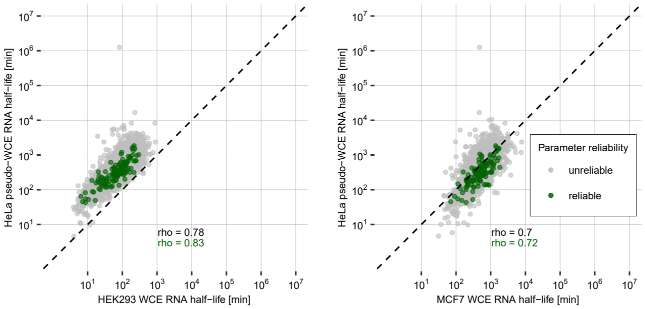


Supplemental Figure S15: RNA half-life comparison between estimates obtained by our two-compartment model and estimates from the literature [37]. For comparison, pseudo-WCE half-lives were calculated by taking the sum of our nuclear and cytosolic half-life estimates.

Further, we also correlated our pseudo-WCE estimates against cytosolic RNA half-lives from Zuckerman et al. (2020) [45] (Supplementary Figure S16).

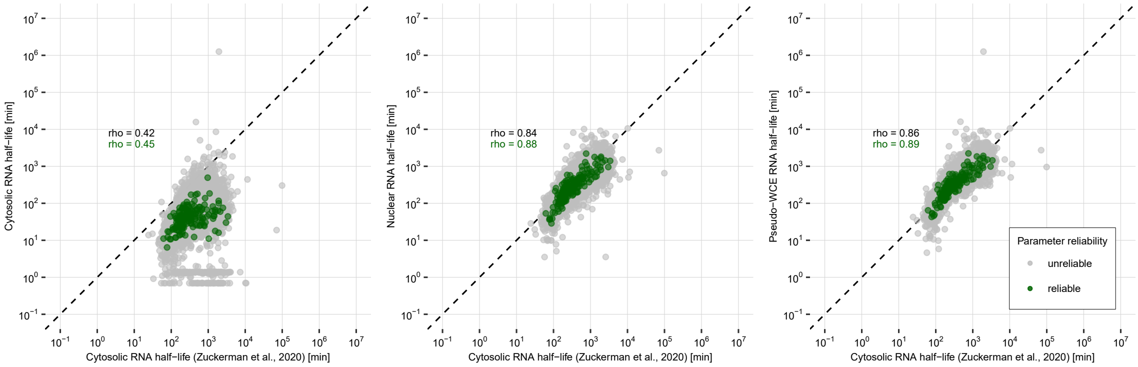


Supplemental Figure S16: Comparison between RNA half-lives from a two-copartment model and estimates obtained by a one-compartment model fit to cytosolic fraction only [45].

#### 5.9 Cytosolic / nuclear RNA ratio estimation

For a set $G$ of transcripts/3'UTRs/peaks considered, let $t_g$,$n_g$, $c_g$ the relative abundances of the total, nuclear and cytosolic RNA population of $g \in G$, respectively (where the abundance is relative to all total/nuclear/cytosolic transcripts that belong to $G$, respectively, i.e., $\sum_{g \in G} t_g = \sum_{g \in G} n_g = \sum_{g \in G} c_g = 1$).

**Spherical median regression.** The set $G$ here is all 3'UTRs with at least 50 counts in the total fraction of control sample (time $t = 0$min). There are 2662 such 3'UTRs, corresponding to a fraction of 4.3% of all annotated 3'UTRs. The selected 3'UTRs account for 84% of all reads assigned to 3'UTRs. If $T$,$N$ and $C$ are numbers that transform total/nuclear/cytosolic relative abundances to (average) molecule numbers per cell, the nuclear and the cytosolic fraction need to sum to the total fraction,

$$Tt_g = Nn_g + Cc_g \quad , \; g \in G$$

This means that $v^T = (T, -N, -C)$ is orthogonal to the plane spanned by the expression vectors $x_g = (t_g, n_g, c_g)^T$, $g \in G$, i.e., $v^T x_g = 0$. The vector $v$ is only determined up to scaling, but this is enough to uniquely determine the nuclear/cytosolic ratio $r$ of the total number of molecules,

$$r = \frac{\sum_g Nn_g}{\sum_g Cc_g} = \frac{N \sum_g n_g}{C \sum_g c_g} = \frac{N}{C}$$

We might therefore perform a linear regression to determine $v$, but this would lead to wrong results, as too many assumptions of linear regression are violated (to name just a few: the data is not drawn from a normal distribution, it is not homoschedastic, there are frequent outliers, linear regression assumes an error in the endpoint and not in the covariates while this is just the other way round here). Thus, we apply a robust, non-parametric regression procedure which resembles the Theil-Sen estimator in standard linear regression: We randomly pick three different transcripts from $G$ and calculate the normal vector of the plane spanned by these three points in $\mathbb{R}^3$. Doing this $R = 10^5$times gives $R$ normal directions represented by unit vectors $v_i$, $i = 1, ..., R$, $\|v_i\| = 1$. Our estimate is then obtained as the geometric median $\bar{v}$ of the $v_i$, where the median is defined as the (a) point minimizig the average distance of the $v_i$, $i = 1, ..., R$, to $\bar{v}$ according to some metric $d$,

$$\bar{v} = \underset{v \in \mathbb{R}^3, \|v\|=1}{\operatorname{argmin}} \frac{1}{R} \sum_{i=1}^{R} d(v_i, v) \tag{31}$$

In Euclidean space, $d(x, y) = \|x - y\|$, and finding $\bar{v}$ in Equation (31) is known as Fermat's problem, and the result $\bar{v}$ is called the geometric median. The resulting loss function is convex, and a fast algorithm to solve the minimization task is a gradient descent procedure known as Weiszfeld algorithm ([63], see [**?**] for a concise treatment). However, since the unit vectors merely define a direction, they are better represented as points in projective space $\mathbb{P}^2 = (\mathbb{R}^3 \setminus \{0\}) / \sim$, where $x \sim y$ if $x = \alpha y$ for some $\alpha \in \mathbb{R} \setminus \{0\}$. Euclidean distance is not a meaningful distance measure in $\mathbb{P}^2$. Therefore, we use the Fubini-Study metric induced by the Euclidean scalar product in $\mathbb{R}^3$. This amounts to measuring the angular distance between (the smaller angle) between two elements $x$ and $y$, $d(x, y) = \arccos \left| \frac{\langle x, y \rangle}{\|x\| \|y\|} \right|$. It can be shown that a similar optmization procedure leads to the desired results for Riemannian manifolds of curvature $\leq 2$, and hence also for $\mathbb{P}^2$ with the Fubini-Study metric [64, 65]. In our case, the geometric median can easily be found using Nelder-Mead optimization (function `optim` in R). The result of the procedure is shown in Main Figure 2E. There, we chose an angular parametrization of the unit vectors to visualize them on a 2d plane: Let $v = (v_1, v_2, v_3) \in \mathbb{R}^3$, $\|v\| = 1$. Then there exists an azimuth angle $\psi \in [0, 2\pi)$ and an altitude angle $\phi \in [0, \pi/2]$ such that $\pm v = (\sin \phi, \cos \phi \cdot \sin \psi, \cos \phi \cdot \cos \psi)$. If $v_1 \neq 0$, this representation is unique. For $v_1 = 0$, we have $\phi = 0$, and if require $\psi \in [0, \pi)$, the choice of $\psi$ is also unique in this case. This defines a map $v \mapsto (\psi, \phi) \in [0, 2\pi) \times [0, \pi/2]$ which serves for visualization. Vectors $v = (T, -N, -C)$ with the same $\frac{cyt}{nuc}$ RNA ratio map to ($\psi = \arctan \frac{N}{C}, \phi$), and contour lines of constant $N/C$ ratio can be added to the plot for convenience.

**Nuclear degradation rate as a function of the $\frac{cyt}{nuc}$ RNA ratio.** Here, set $G$ consists of the 251 3'UTRs for which cytosolic degradation rate $\lambda_g$ and the nuclear removal rate $\tau_g + \nu_g$ could be estimated reliably according to our quality criteria (Methods 4.6). In steady state, by Equation (3) yields

$$\frac{Cc_g}{Nn_g} = \frac{\tau_g}{\lambda_g}$$

Multiplying by $\lambda_g$ and adding $\nu_g$ on both sides leads to

$$\frac{C}{N} \cdot \frac{c_g}{n_g} \lambda_g + \nu_g = \tau_g + \nu_g$$

Substituting the nuclear/cytosolic RNA ratio $r = N/C$, we obtain an estimator of the nuclear degradation rate, $\hat{\nu}_g(r)$, as a function of $r$:

$$\hat{\nu}_g(r) = (\tau_g + \nu_g) - r^{-1} \frac{c_g}{n_g} \lambda_g \tag{32}$$

This allows us to calculate the tail probabilities of the empirical distribution of $\{\hat{\nu}_g(r); g \in G\}$,

$$P(\hat{\nu}(r) > 0) = \frac{|\{g \in G \mid \hat{\nu}_g(r) > 0\}|}{|G|}$$

as a function of $r$. As $\nu_g \geq 0$ for all $g$, we expect at least 50% of the estimates to be non-negative. According to Main Figure 2F, this is only the case for $r \geq 4.6$. For ratios $r \leq 1$, less than 6% of all nuclear degradation rate estimates would be positive.

**Spike-in normalisation.** Spike-ins containing constant, well-defined amounts of unique non-host RNAs were added to the nuclear and cytosolic fractions. Let $G$ all 3'UTRs with at least 1 read count in each time point measurement of the time series, plus the set of all spike-in RNAs. For each RNA sample, we calculated a robust average $\bar{n}$ of the relative abundances for the nucleus, $n_h$, $h \in H$, as the interquartile mean, i.e. the mean of the central 50% of all relative expression values lying between the 25% and the 75% quantile of the data. Similarly, $\bar{c}$ was defined as the interquartile mean of $c_h$, $h \in H$. Let $\bar{n}_{\text{spike}}$ respectively $\bar{c}_{\text{spike}}$ the mean expression values of the spike-in RNAs we used in the nucleus respectively in the cytosol. As above, let $N$ respectively $C$ the constants that transfer relative abundances in the nucleus respectively the cytosol to molecules per cell. As the total amount of the spike-ins that has been added to each fraction is constant. It can be calculated as $N\bar{n}_{\text{spike}} = C\bar{c}_{\text{spike}}$. An estimate of the $\frac{cyt}{nuc}$ RNA ratio is therefore obtained by

$$\hat{r} = \frac{C\bar{c}}{N\bar{n}} = \frac{\bar{c}}{\bar{n}} \cdot \frac{C\bar{c}_{\text{spike}}}{N\bar{n}_{\text{spike}}} \cdot \frac{\bar{n}_{\text{spike}}}{\bar{c}_{\text{spike}}} = \frac{\bar{c}}{\bar{n}} \cdot \frac{\bar{n}_{\text{spike}}}{\bar{c}_{\text{spike}}}$$

#### 5.10 Retrieval of gene-specific features

GENCODE annotation was used to retrieve information about exon counts and lengths, as well as CDS lengths. If a gene encodes for multiple transcript isoforms, these lengths and counts were averaged. We retrieved information about 3'UTR lengths from our UTR annotation file described in Methods 4.3. Information about a transcript's GC content and length was retrieved from Ensembl biomart. We always considered the longest transcript isoform of a gene for comparison, assuming this transcript to be the dominantly expressed one. For each 3'UTR we measured in our experiments, we associated the corresponding half-life estimates with the gene-specific features via their gene names. To avoid ambiguity, 3'UTRs with multiple gene annotations were removed from the analyses.

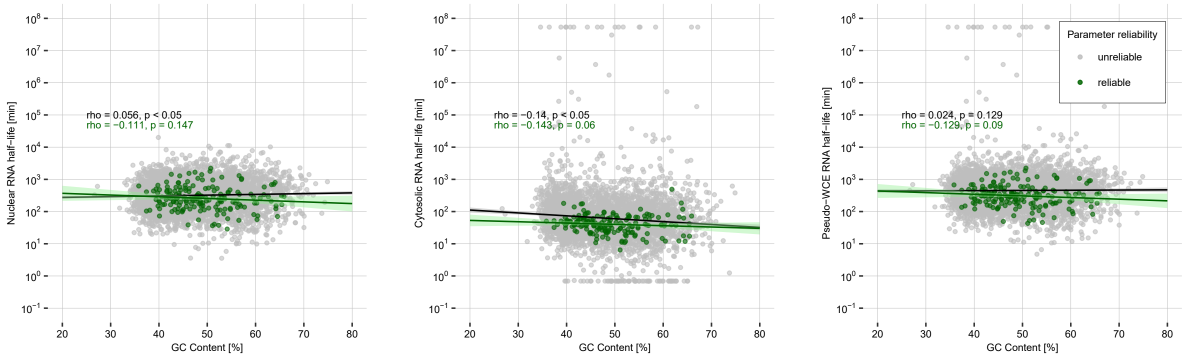


Supplemental Figure S17: Scatterplot comparing the half-life estimates of our two-compartment model with the corresponding GC content of a respective gene.

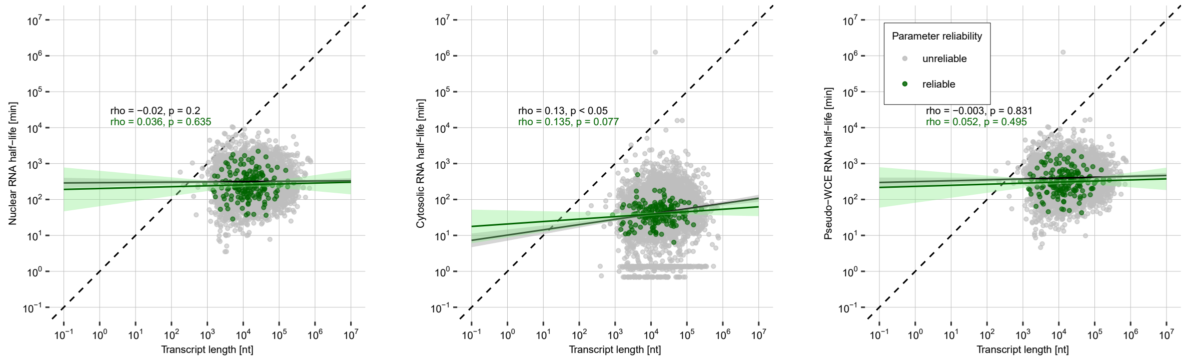


Supplemental Figure S18: Scatterplot comparing the half-life estimates of our two-compartment model with the corresponding transcript length of a respective gene.

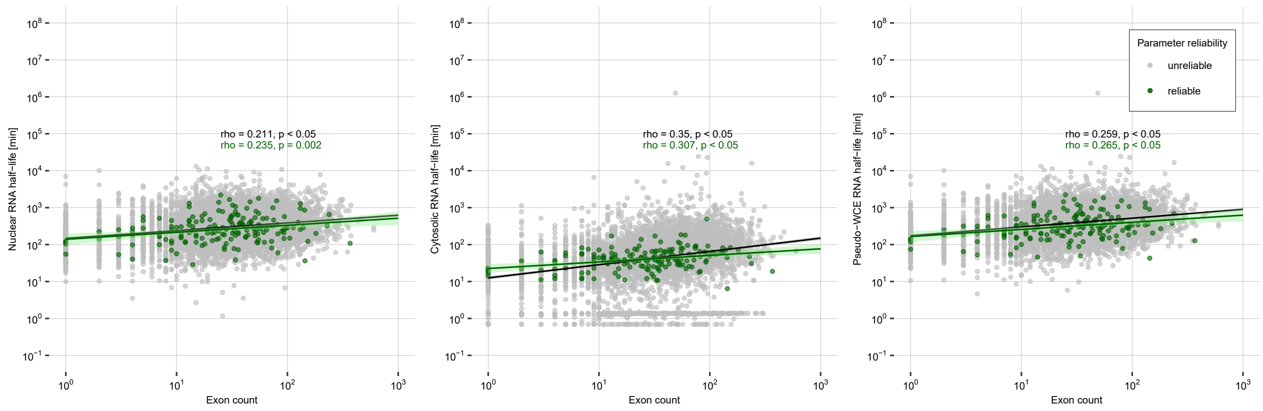


Supplemental Figure S19: Scatterplot comparing the half-life estimates of our two-compartment model with the corresponding exon count of a respective gene.

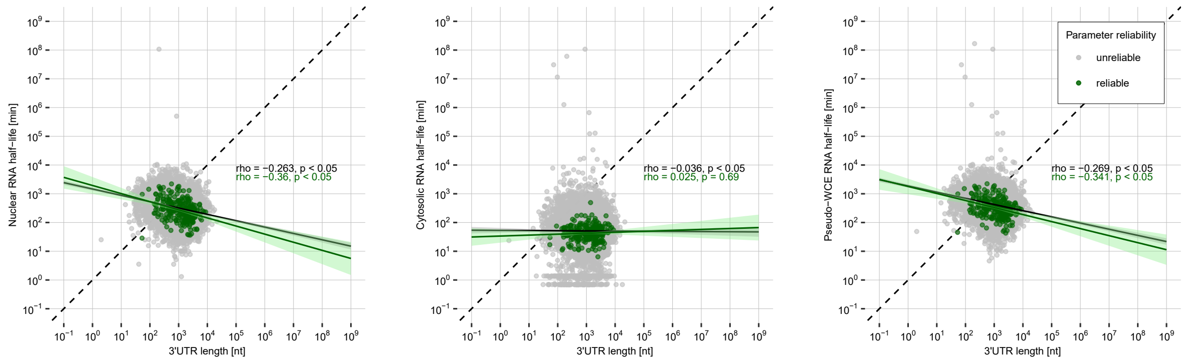


Supplemental Figure S20: Scatterplot comparing the half-life estimates of our two-compartment model with the corresponding 3'UTR length of a respective gene.

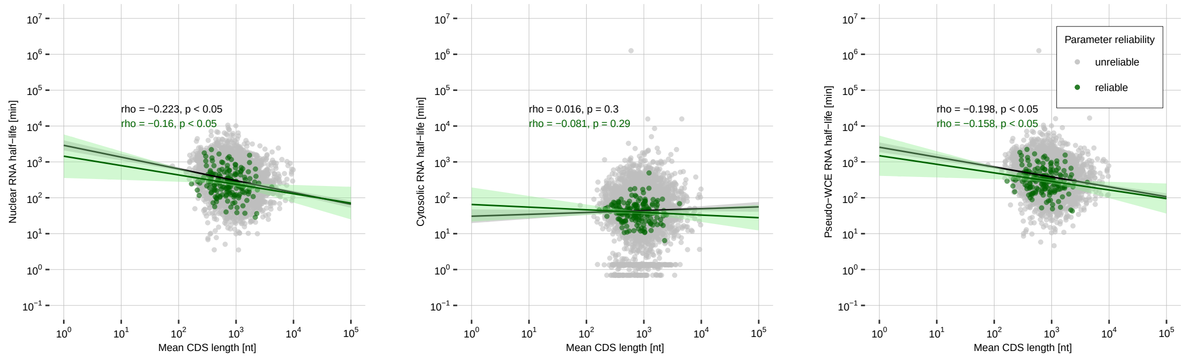


Supplemental Figure S21: Scatterplot comparing the half-life estimates of our two-compartment model with the corresponding CDS length of a respective gene.

#### 5.11 Difference in RNA half-lives between 3'UTR isoforms

We used peak calling to define potential 3'UTR isoforms in an intragenic region. To that end, we defined peaks located in the same 3'UTR but separated by at least 100 nucleotides to be different 3'UTR isoforms. Differences in nuclear respectively cytosolic half-lives between these isoforms are shown in Supplemental Figure SS22-S23.

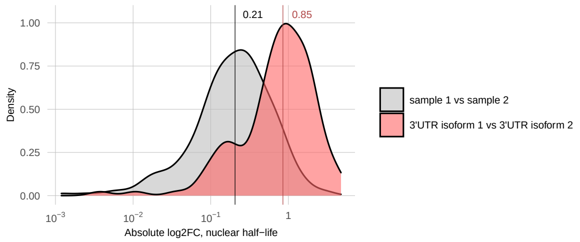


Supplemental Figure S22: Difference in nuclear half-lives of potential 3'UTR isoforms. Some 3'UTRs harbored at least two sufficiently covered and distant peaks, representing potential 3'UTR isoforms (242 peaks from 118 3'UTRs). The absolute log fold change of both nuclear and cytosolic half lifes was computed for all pairwise comparisons of the potential isoform irrespective of the half life estimate reliability. As comparison, absolute log2 fold changes were calculated for an isoform's estimates between time series 1 and 2. The medians are indicated by the red and black lines with corresponding text labels.

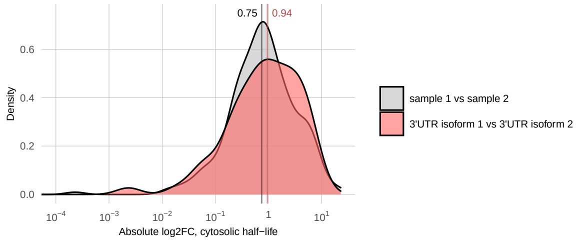


Supplemental Figure S23: Difference in cytosolic half-lives of potential 3'UTR isoforms. Some 3'UTRs harbored at least two sufficiently covered and distant peaks, representing potential 3'UTR isoforms (242 peaks from 118 3'UTRs). The absolute log fold change of both nuclear and cytosolic half lifes was computed for all pairwise comparisons of the potential isoform irrespective of the half life estimate reliability. As comparison, absolute log2 fold changes were calculated for an isoform's estimates between time series 1 and 2. The medians are indicated by the red and black lines with corresponding text labels.

#### 5.12 RBP analysis

Protein-RNA-binding information was obtained from an eCLIP experiment from the ENCODE project [43, 42]. The data set comprises 120 distinct RBP-RNA interaction profiles in K562 cells. We considered BED files for the hg19 genome assembly only, to match the genome assembly used in our SLAM-seq experiments. We used GENCODE annotation [66] for the mapping of protein binding sites to genes. Based on the fact that poly-adenylated mRNA barely contain any introns, we excluded intronic protein binding sites. To control the number of false positives, we only kept binding sites passing stringent cutoffs (-log10(p-value) $\geq$ 5 and log2(fold-enrichment) $\geq$ 3).

Considering only 3'UTRs with a reliable rate estimate, we define a 3'UTR to be bound by an RBP, if this region can be uniquely assigned to one gene, and this gene also contains a binding site for that RBP. This way we obtain, for each RBP, a set of bound and a set of unbound transcripts. A Wilcoxon rank sum test was applied to compare if the nuclear or cytosolic half-life distribution of the bound transcripts differs significantly from the one of the unbound transcripts. The p-values were Bonferroni corrected for multiple testing.

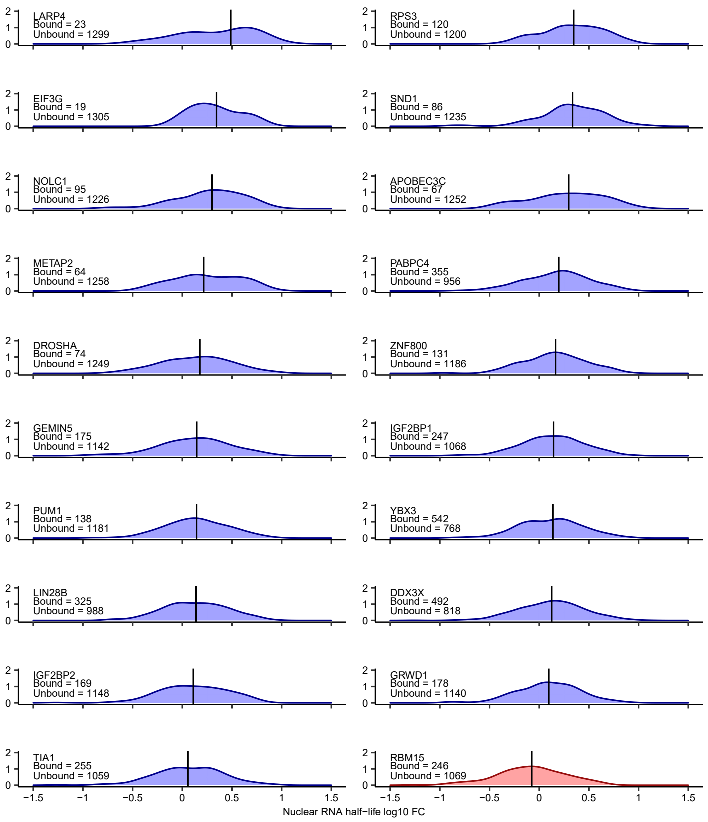


Supplemental Figure S24: Density plots for distinct RBPs from the eCLIP analysis. Shown are the distributions of the nuclear half-lives from RNAs putatively bound to the respective RBP according to eCLIP experiments. Log10 fold-changes were calculated by taking the log10 of the nuclear half-lives and substracting the log10 of the global median (e.g. the median of all bound and unbound transcript half-lives) from these values. Blue densities indicate distributions with a higher median half live of the bound transcripts as the global median. The red density indicates a shorter median transcript half-life of the bound RNAs in comparison with the global median. Black bars indicate the median of the respective distribution.

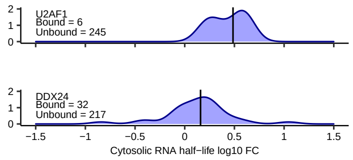


Supplemental Figure S25: Density plots for distinct RBPs from the eCLIP analysis. Shown are the distributions of the cytosolic half-lives from RNAs putatively bound to the respective RBP according to eCLIP experiments. Log10 fold-changes were calculated by taking the log10 of the cytosolic half-lives and substracting the log10 of the global median (e.g. the median of all bound and unbound transcript half-lives) from these values. Blue densities indicate distributions with a higher median half live of the bound transcripts as the global median. The red density indicates a shorter median transcript half-life of the bound RNAs in comparison with the global median. Black bars indicate the median of the respective distribution.

#### 5.13 Assessment of half-lives of lncRNAs and mRNAs translated at the ER

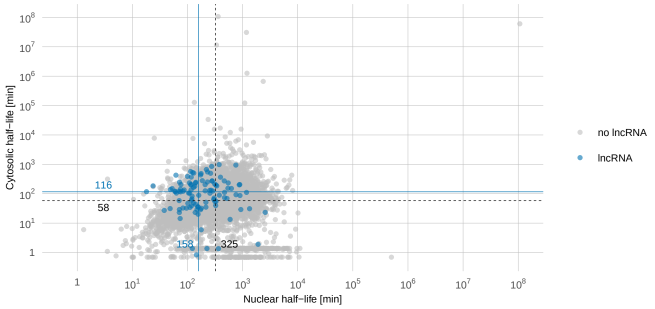


Supplemental Figure S26: Half-life estimates of lncRNAs in comparison to all 3'UTRs. From all sufficiently covered 3'UTRs, some were annotated as lncRNAs (blue data points). The blue lines indicate the median half-lives of the lncRNAs, the black lines indicate the median half-lives of all sufficiently covered 3'UTRs, respectively.

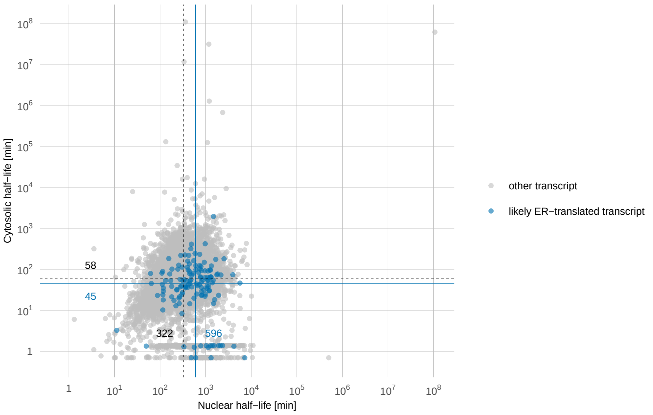


Supplemental Figure S27: Median half lifes of transcripts likely translated at the ER. From all expressed 3'UTRs, two subsets of transcripts were selected which are suspicious of being translated at the ER using appropriate GO. The blue and black lines with corresponding text labels indicate the median half lifes of the 3'UTR subsets and all sufficiently covered 3'UTRs, respectively.

#### 5.14 Model including nuclear retention

In comparison to our standard two-compartment model (Eqs 1 and 2), we split the nuclear RNA $N$ into two fractions, namely the fraction $E$ that will be exported, and the retained fraction $R$, $N = E + R$. We assume that the decision whether an RNA is exported or not is made co-transcriptionally. Hence we introduce an additional retention paramter $r \in [0, 1]$ which determines the probability by which a transcript will be retained after completion of the transcription process.This leads to the ODE system

$$\dot{E} = \mu(1 - r) - (\nu + \tau)E \tag{33}$$

$$\dot{C} = \tau E - \lambda C \tag{34}$$

$$\dot{R} = \mu r - \nu R \tag{35}$$

with the boundary conditions $R_{new}(0) = E_{new}(0) = C_{new}(0) = 0$. When cells are in dynamic steady state, we have $R_\infty = \frac{\mu r}{\nu}$, $E_\infty = \frac{\mu(1-r)}{\nu+\tau}$, $C_\infty = E_\infty \frac{\tau}{\lambda}$. The ODE for old RNA equal those for new, with the exception that the terms including $\mu$ are omitted (or, $\mu$ is set to 0). This leads to the closed form solutions

$$\begin{aligned} R_{old}(t) &= R_\infty \cdot e^{-\nu t} && (36) \\ E_{old}(t) &= E_\infty \cdot e^{-(\nu+\tau)t} && (37) \\ C_{old}(t) &= C_\infty \cdot (\frac{\lambda e^{-(\nu+\tau)t} - (\nu+\tau)e^{-\lambda t}}{\lambda - (\nu+\tau)}) && (38) \end{aligned}$$

Correspondingly, the new fraction time curves are obtained by subtracting the old RNA levels from the steady state levels:

$$\begin{aligned} R_{new}(t) &= R_\infty \cdot (1 - e^{-\nu t}) && (39) \\ E_{new}(t) &= E_\infty \cdot (1 - e^{-(\nu+\tau)t}) && (40) \\ C_{new}(t) &= C_\infty \cdot \left(1 - \frac{\lambda e^{-(\nu+\tau)t} - (\nu+\tau)e^{-\lambda t}}{\lambda - (\nu+\tau)}\right) && (41) \end{aligned}$$

Subsequently, we get nuclear, respectively cytsolic new/total ratios for each timepoint t by:

$$\frac{N_{new}(t)}{N_\infty} = \frac{R_{new}(t) + E_{new}(t)}{R_\infty + E_\infty} = \frac{\frac{r}{\nu} \cdot (1 - e^{\nu t}) + \frac{1-r}{\nu+\tau} \cdot (1 - e^{-(\nu+\tau)t})}{\frac{r}{\nu} + \frac{1-r}{\nu+\tau}} \quad (42)$$

$$\frac{C_{new}(t)}{C_\infty} = 1 - \frac{\lambda e^{-(\nu+\tau)t} - (\nu+\tau)e^{-\lambda t}}{\lambda - (\nu+\tau)} \quad (43)$$

Given the observed nuclear and cytosolic new/total RNA ratios, Equations (42), (43) are then used to fit the parameters $\tau, \lambda, r$, for a grid of values $\nu$. We varied the value for the nuclear degradation rate $\nu$ extensively, to ensure realistic levels of accumulation of the retained RNA fraction. We used a three-dimensional MCMC to obtain a representative sample and an optimized fit for the nuclear export rate $\tau$,the nuclear retention rate $r$ and the cytosolic degradation rate $\lambda$. The sampler was initialized with the results of a global optimization algorithm (Differential Evolution algorithm, R command DEoptim). Next, the component-wise medians of the MCMC samples were computed after burn-in. These medians were used for another global optimization (Differential Evolution algorithm) to obtain the final parameter estimates for $\tau, r$ and $\lambda$. Parameter fitting was seperaterly perforemd for both samples, and estimates averaged afterwards. Further, we calculate the percentage of retained transcripts as

$$\frac{R_\infty}{R_\infty + N_\infty} = \frac{\tau r + \nu r}{\tau r + \nu} \quad (44)$$

#### 5.15 One-compartment model of the cytosolic RNA

A one-compartment model of the cytosolic compartment is given by

$$\dot{C}(t) = \mu - \lambda C \quad (45)$$

with boundary conditions $C_{new}(0) = 0$ and $C_{old}(0) = C_\infty = \frac{\mu}{\lambda}$. Fitting of $\lambda$ was performed as described before for the nuclear compartment (Methods 4.5), separately for the two time series, and estimates were averaged afterwards.

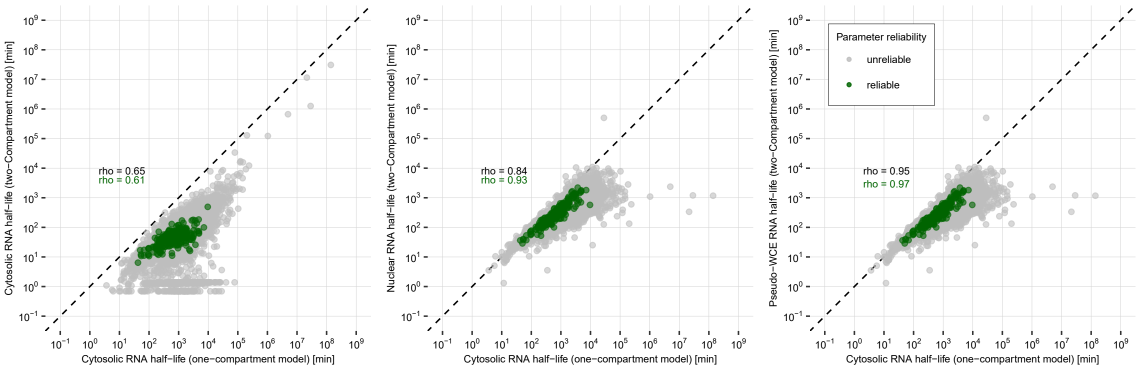


Supplemental Figure S28: Comparison of estimates derived from a two-compartment and one-comparmtent model. For visualisation purposes, the plot was cropped leaving out 2 data points.

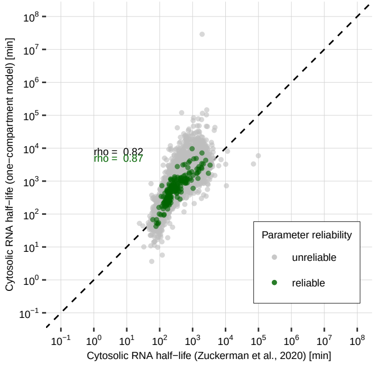


Supplemental Figure S29: Correlation of a simple exponential cytosolic decay model fit results with cytosilc half-lives obtained through SLAM-seq on the cytosolic fraction of MCF7 cells by Zuckerman et al. (2020) [45]. gray dots represent expressed 3'UTRs and green dots portray 3'UTRs that passed our reliability criteria for the cytosolic compartment.

#### 5.16 T>C conversion and mismatch statistics

Conversions originating from the labeling are expected to show as A>G on the (-) and T>C on the (+) strand. Expectedly and reassuringly, the labeling conversion rates clearly show an increasing trend over time, while the other rates stay constant and low. Also, the cytosolic rates increase less less and slower than the nuclear rates, as the export of newly synthesized, labeled RNA transcripts from the nucleus to the cytosol requires time.

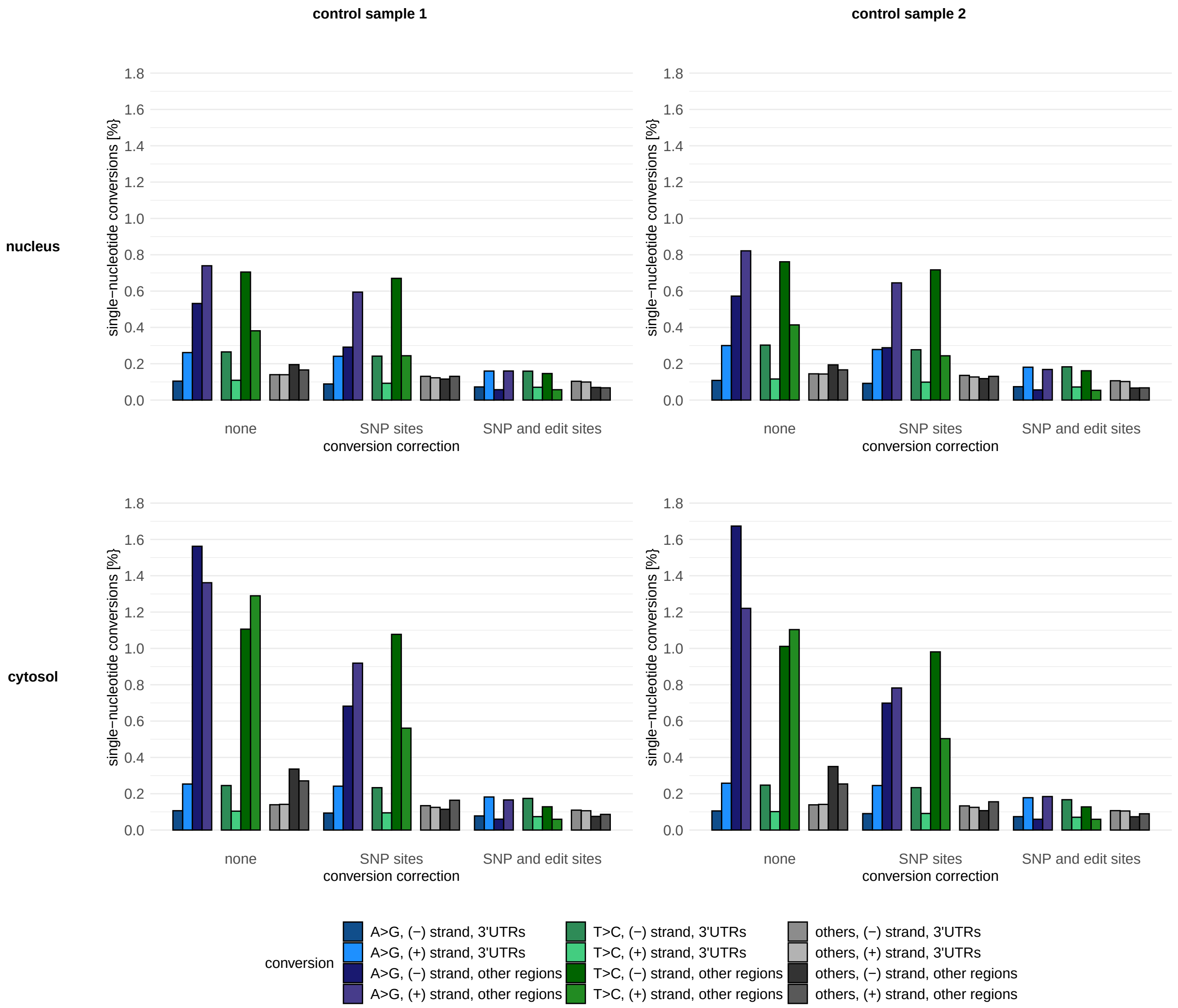


Supplemental Figure S30: Conversion statistics of the control samples. No conversion correction: mismatch rates as observed in the primary alignment. SNP-sites correction: mismatch rates as observed in the primary alignment when excluding all nucleotide positions annotated as known SNP site. SNP and edit-site correction: conversions as observed in the primary alignment when excluding all nucleotide positions annotated as known SNP site or tagged as potential editing sites. A>G (T>C) conversions: A>G (T>C) conversions as observed in the read alignment. Other conversions: average of all non-A>G and non-T>C conversions observed in the read alignment. Conversions of (-) strand ((+) strand) 3'UTRs: conversions of all uniquely mapped reads which are assigned to an annotated (-) strand ((+) strand) 3'UTR. Conversions of other regions on the (-) strand ((+) strand): conversions of all uniquely mapped reads which are assigned to a non-3'UTR peak on the (-) strand ((+) strand) with a minimum coverage of 5 in the sample and compartment.

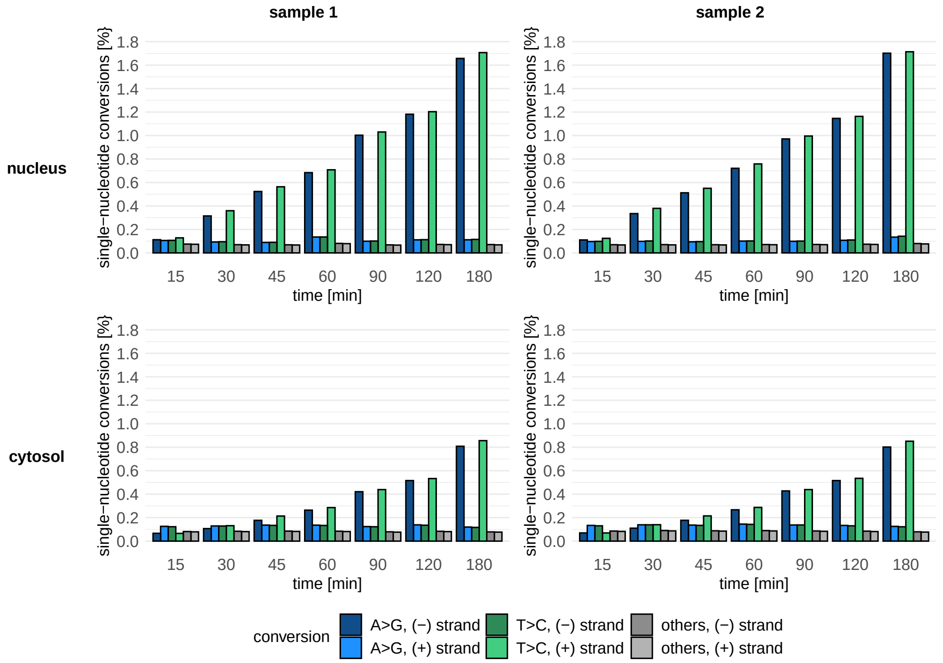


Supplemental Figure S31: Conversion statistics of the SLAM-seq labeling samples, shown for the reads assigned to annotated 3'UTRs. Statistics are corrected for both annotated SNP sites and potential editing sites. A>G (T>C) conversions: A>G (T>C) conversions as observed in the read alignment. Other conversions: average of all non-A>G and non-T>C conversions observed in the read alignment. Conversions of the EM algorithm(-) strand ((+) strand): conversions of all uniquely mapped reads which are assigned to an annotated (-) strand ((+) strand) 3'UTR.

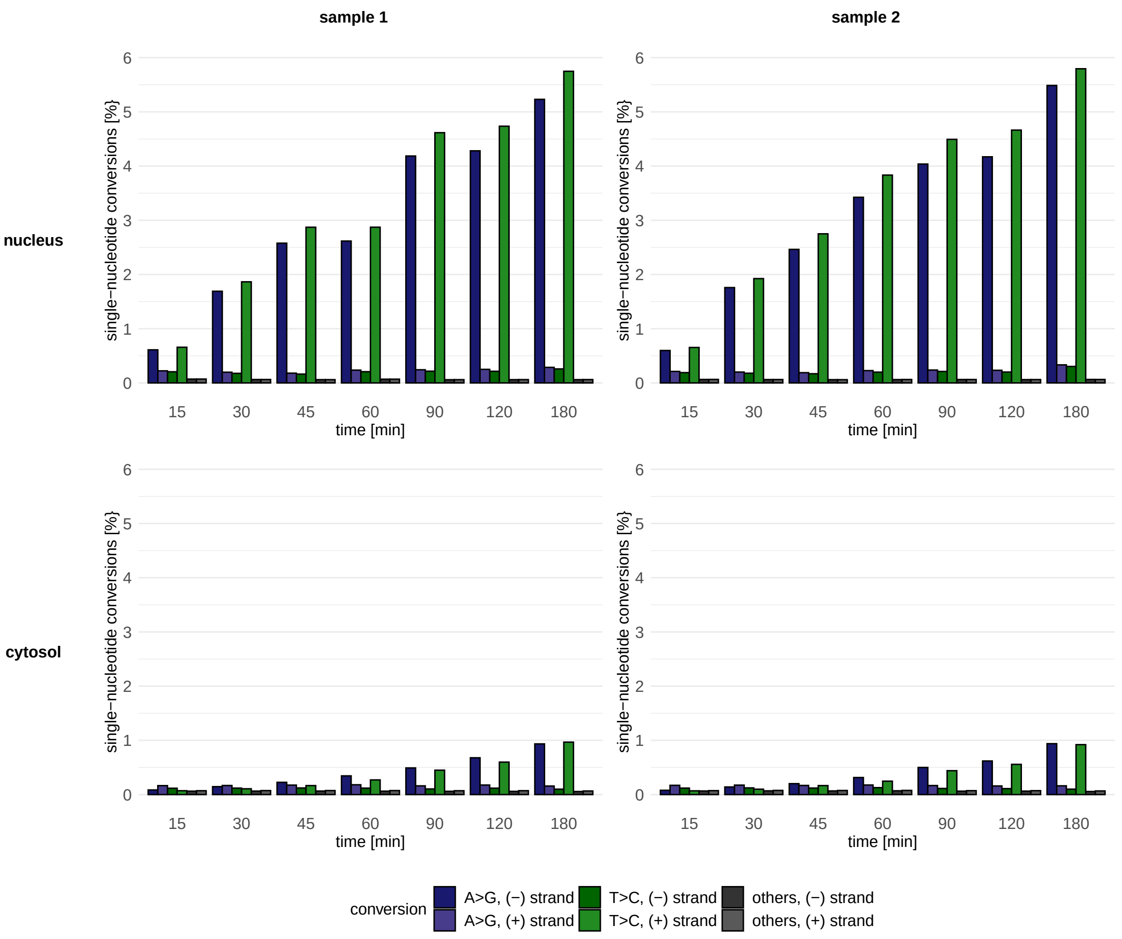


Supplemental Figure S32: Conversion statistics of the SLAM-seq labeling samples, shown for the reads assigned to non-3'UTR peaks. For each measurement, peaks with a coverage of less than 5 in that measurement were excluded from the calculations. Statistics are corrected for both annotated SNP sites and potential editing sites. A>G (T>C) conversions: A>G (T>C) conversions as observed in the read alignment. Other conversions: average of all non-A>G and non-T>C conversions observed in the read alignment. Conversions of the (-) strand ((+) strand): conversions of all uniquely mapped reads which are assigned to an annotated (-) strand ((+) strand) non-3'UTR peak.
